# Amino acid transporter SLC38A5 regulates developmental and pathological retinal angiogenesis

**DOI:** 10.1101/2021.09.01.458523

**Authors:** Zhongxiao Wang, Felix Yemanyi, Shuo Huang, Chi-Hsiu Liu, William R. Britton, Steve S. Cho, Alexandra K. Blomfield, Yohei Tomita, Zhongjie Fu, Jian-Xing Ma, Wen-Hong Li, Jing Chen

## Abstract

Amino acid metabolism in vascular endothelium is important for sprouting angiogenesis. SLC38A5 (solute carrier family 38 member 5), an amino acid (AA) transporter, shuttles neutral AAs across cell membrane, including glutamine, which may serve as metabolic fuel for proliferating endothelial cells (ECs) to promote angiogenesis. Here we found that *Slc38a5* is highly enriched in normal retinal vascular endothelium, and more specifically in pathological sprouting neovessels. *Slc38a5* is suppressed in retinal blood vessels from *Lrp5^-/-^* and *Ndp^y/-^* mice, both genetic models of defective retinal vascular development with Wnt signaling mutations. Additionally, *Slc38a5* transcription is directly regulated by Wnt/β-catenin signaling. Genetic deficiency of *Slc38a5* in mice substantially delays retinal vascular development and suppresses pathological neovascularization in oxygen-induced retinopathy modeling ischemic proliferative retinopathies. Inhibition of *SLC38A5* in retinal vascular ECs impairs EC proliferation and angiogenic function, suppresses glutamine uptake, and dampens vascular endothelial growth factor receptor 2 (VEGFR2). Together these findings suggest that SLC38A5 is a new metabolic regulator of retinal angiogenesis by controlling AA nutrient uptake and homeostasis in ECs.

**Significance Statement:** Amino acid metabolism in vascular endothelium is important for angiogenesis. SLC38A5 (solute carrier family 38 member 5) is an amino acid (AA) transporter for shuttling neutral AAs such as glutamine across cell membrane. Our work demonstrate that *Slc38a5* is highly enriched in retinal vascular endothelium. SLC38A5 regulates endothelial cell glutamine uptake and vascular growth factor receptors to impact blood vessels growth in retinal development and in retinopathies. This work uncovered a novel role of SLC38A5 as a metabolic regulator of retinal angiogenesis by controlling AA nutrient uptake and homeostasis in blood vessel endothelium. Findings from this study also suggest that targeting SLC38A5 or relevant AAs can be a new way to protect against retinopathy.

## Introduction

Angiogenesis, the growth of new blood vessels from existing vessels, is important in both development and disease (1). In the developing eye, formation of blood vessels allows delivery of nutrients and removal of metabolic waste from neuronal retinas (2, 3). In vascular eye diseases, specifically proliferative retinopathies, such as retinopathy of prematurity and diabetic retinopathy, abnormal proliferation of pathological blood vessels may lead to retinal detachment and vision loss (4–6). Angiogenesis is coordinated by many pro-angiogenic factors, such as vascular endothelial growth factor (VEGF), and angiogenic inhibitors. The balance of these factors maintains vascular endothelium in proper homeostasis. In addition to growth factors, metabolism in vascular endothelial cells (ECs) has been recognized as a driving force of angiogenesis (7, 8). Both glucose glycolysis (9–11) and fatty acid oxidation (12–15) play major roles in regulating angiogenesis, and more specifically in ocular angiogenesis (16–18). Moreover, metabolism of amino acids (AAs), such as glutamine, is increasingly established as an essential energy source in EC sprouting and angiogenesis (7, 19).

Several AAs may serve as metabolic fuel for proliferating ECs to promote angiogenesis, including glutamine, asparagine, serine and glycine (20–23). Glutamine, the most abundant non-essential amino acid in the body, is a key carbon and nitrogen source and can be metabolized to sustain EC growth (20, 24). It can act as an anaplerotic source of carbon to replenish the tricarboxylic acid (TCA) cycle and support protein and nucleotide synthesis in EC growth (20, 21). Glutamine deprivation severely impairs EC proliferation and vessel sprouting, and pharmacological blockade of glutaminase 1, an enzyme that converts glutamine to glutamate, inhibits pathological angiogenesis (20). In addition to anaplerosis, glutamine also regulates angiogenesis by mediating amino acid synthesis, functioning as a precursor to other amino acids such as glutamic acid, aspartic acid, and asparagine, as well as mediating macromolecule synthesis and redox homeostasis (20, 21, 25–28). When glutamine is low, asparagine may be used as an alternative AA source (29), and can partially rescue glutamine-restricted EC defects (20, 21). In brain vascular ECs with barrier polarity, glutamine transport relies on facilitative transport systems on the luminal membrane of EC and sodium-dependent transport systems on the EC abluminal membrane, to actively transport glutamine from extracellular environment into ECs (30).

Solute carrier family 38 member 5 (SLC38A5, also known as SNAT5: system N sodium-coupled amino acid transporter 5) transports neutral AAs across cell membrane, including glutamine, asparagine, histidine, serine, alanine, and glycine (31). Previously, SLC38A5 was found to mediate transcellular transport of AAs in brain glial cells (32). SLC38A5 is also a recently identified marker for pancreatic progenitors (33), where it is important for L-glutamine-dependent nutrient sensing and pancreatic alpha cell proliferation and hyperplasia (34, 35). In the eye, SLC38A5 was previously found in Müller glial cells and retinal ganglion cells (36, 37), yet its localization and function in other retinal cells, including vascular ECs, are less clear. We and others previously found that *Slc38a5* was drastically down-regulated in both *Lrp5^-/-^* and *Ndp^y/-^* retinas (38–40), experimental models of two genetic vascular eye diseases: familial exudative vitreoretinopathy (FEVR) and Norrie disease respectively. Both disease models have genetic mutations in Wnt signaling and share similar retinal vascular defects including initial incomplete or delayed vascular development, absence of deep retinal vascular layer, followed by a secondary hypoxia-driven increase in VEGF in the retina, and tuft-like neovascularization in the superficial layer of the retinal vasculature (41–44). *Slc38a5* is down-regulated 7-10 fold in *Lrp5^-/-^* retinas and *Ndp^y/-^* retinas during development (38, 40), indicating a potential strong connection of *Slc38a5* with retinal blood vessel formation.

This study explored the regulatory functions of SLC38A5 in retinal angiogenesis during development and in disease. We found that *Slc38a5* expression is enriched in retinal blood vessels and its transcription is directly regulated by Wnt signaling. Moreover, genetic deficiency of *Slc38a5* impairs both developmental retinal angiogenesis and pathological retinal angiogenesis in a mouse model of oxygen-induced retinopathy (OIR), modeling proliferative retinopathies. Furthermore, inhibition of SLC38A5 decreases EC angiogenic function *in vitro,* and dampens EC glutamine uptake and growth factor signaling including VEGFR2. Together these findings identified a pro-angiogenic role of SLC38A5 in ocular angiogenesis, and suggest this transporter as a potential new target for developing therapeutics to treat pathological retinal angiogenesis.

## Methods and materials

### Animals

All animal studies described in this paper were approved by the Boston Children’s Hospital Institutional Animal Care and Use Committee (IACUC), and also adhered to the ARVO Statement for the Use of Animals in Ophthalmic and Vision Research. *Slc38a5^-/-^* mice (100% of C57BL/6 background) were generated and knockout validated previously (35). Both male and female *Slc38a5* knockout mice were used in experiments and littermate WT controls from the same breeding colony were used as comparison. C57BL/6J mice were obtained from Jackson Laboratory (stock no: 000664) and were used for siRNA treatment and laser capture microdissection experiments as well as wild type (WT) control mice for *Lrp5^-/-^* (stock no. 005823). *Ndp^y/-^* (stock no. 012287) were also obtained from the Jackson Laboratory (Bar Harbor, ME). Male *Ndp^y/+^* mice were used as control for *Ndp^y/-^* for X-linked *Ndp* gene.

### Oxygen-induced retinopathy (OIR)

The OIR mouse model was performed as previously described (45, 46). Newborn mouse pups with nursing mothers were exposed to 75 ± 2 % oxygen from postnatal day (P) 7 to P12 and returned to room air until P17. Mice were anesthetized with ketamine/xylazine and euthanized by cervical dislocation, followed by retinal dissection and blood vessel staining.

### Intravitreal injection of siRNA

Intravitreal injections were performed in C57BL/6J mouse pups at various developmental time points or with OIR, according to previously established protocols (47–50). Briefly, mice were anesthetized with isoflurane in oxygen. 1 μg of si-Slc38a5 (ThermoFisher, Cat# 4390771) dissolved in 0.5μL of vehicle solution was injected using a 33-gauge needle behind the limbus of the left eye, whereas the contralateral right eye of the same animal was injected with an equal amount of negative control scrambled siRNA (si-Ctrl) (ThermoFisher, Cat# 4390844). After injection, eyes were lubricated with sterile saline and an antibiotic eye ointment was applied. At specified days after injection, mice were sacrificed, and retinal vasculature was analyzed.

### Retinal dissection and vessel staining

Mouse pups at various developmental time points or post-OIR exposure were sacrificed, retinas dissected, and blood vessels stained according to previous protocols (46). Isolated eyes were fixed in 4% paraformaldehyde in phosphate-buffered saline (PBS) for 1 hour at room temperature. Retinas were dissected, stained with labeled *Griffonia Simplicifolia* Isolectin B4 (Alexa Fluor 594 conjugated, Cat# 121413; Invitrogen), and flat-mounted onto microscope slides (Superfrost/Plus, 12-550-15; Thermo Fisher Scientific, Waltham, MA) with photoreceptor side down and embedded in antifade reagent (SlowFade, S2828; Invitrogen). Retinas were imaged with a fluorescence microscope (AxioObserver.Z1 microscope; Carl Zeiss Microscopy) and images were merged to cover the whole flat-mounted retina. For imaging deeper retinal layers, fixed retinas were permeabilized first with PBS in 1% Triton X-100 for 30 min, followed by isolectin B4 staining with 0.1% TritonX-100 in similar processes as described above.

### Quantification of retinal vascular development, vaso-obliteration and pathological neovascularization in OIR

Quantification of developmental vasculature were performed by using Adobe Photoshop (Adobe Systems, San Jose, CA, USA) and ImageJ from NIH according to previous protocols (3, 46). Retinal vascular areas were expressed as a percentage of the total retinal areas. n is the number of retinas quantified.

Quantification of retinal vaso-obliteration and neovascularization in OIR retinas followed previously described protocols (51–53) by using Adobe Photoshop and ImageJ. The avascular area absent of isolectin B4 staining was outlined and calculated as a percentage of the whole retinal area. Pathological neovascularization was recognized by the abnormal aggregated morphology and was quantified using a computer-aided SWIFT_NV method and also normalized as percentage of the whole retina area (51). Quantification was done with the identity of the samples masked.

### Laser capture microdissection of retinal vessels

Laser capture micro-dissection (LCM) of retinal vessels were carried out based on previous protocols (48, 54, 55). Mouse eyes were embedded in optimal cutting temperature (OCT) compound, sectioned at 10 μm, and mounted on polyethylene naphthalate glass slides (Cat# 115005189, Leica). Frozen sections were dehydrated, briefly washed, then stained with isolectin B4 to visualize blood vessels. Retinal blood vessels were laser capture microdissected with Leica LMD 6000 system (Leica Microsystems). Micro-dissected samples were collected in lysis buffer from the RNeasy micro kit (Cat# 74004, Qiagen, Chatsworth, MD, USA), followed by RNA isolation and RT-qPCR.

### Single-cell transcriptome analysis

Gene expression of *Slc38a5* and endothelial cell marker *Pecam1* in mouse retinal cells types was identified using the online single-cell dataset: Study - P14 C57BL/6J mouse retinas (https://singlecell.broadinstitute.org/single_cell/study/SCP301) (56). Similarly, gene expression of *SLC38A5* and *PECAM1* was identified in human retinal single cell dataset - Cell atlas of the human fovea and peripheral retina (https://singlecell.broadinstitute.org/single_cell/study/SCP839) (57). Both studies are accessed from Single Cell Portal, Broad Institute. Dot plots of gene expression for different retinal cell types were grouped and displayed.

### EC cell culture and assays of angiogenic function and glutamine uptake

Human retinal microvascular endothelial cells (HRMECs, Cat# ACBRI 181, Cell system) were cultured in completed endothelial culture medium supplemented with culture boost-R (55). Cell between passage 4 to 7 were transfected with *si-SLC38A5* siRNA (Cat# 4392420, Thermo Fisher Scientific) or negative control siRNA (si-Ctrl, Cat # AM4611, Thermo Fisher Scientific). siRNA knockdown was confirmed by RT-qPCR and Western blot of cells collected 48-72 hours after transfection.

HRMEC viability and/or proliferation was assessed at 48 and 72 hours after transfection with *si-SLC38A5* or si-Ctrl using a MTT cell metabolic activity assay kit (Cat# V13154, Life Technologies) as described previously (49). Briefly, HRMECs were incubated for 4 hrs in solutions containing a yellow tetrazolium salt MTT (3-(4,5-dimethylthiazol-2-yl)-2,5-diphenyltetrazolium bromide). NAD(P)H-dependent oxidoreductase in metabolically active live HRMEC reduces MTT to purple formazan crystals. Afterwards a solution containing SDS-HCL was added which dissolve the purple formazan, followed by measurement of absorbance at 570 nm using a multi-well spectrophotometer. Measurement of MTT cellular metabolic activity is indicative of cell viability, proliferation and cytotoxicity. HRMEC migration and tube formation assays were carried out according to previous protocols (48).

Glutamine uptake by HRMEC was performed using a glutamine/glutamate-Glo bioluminescent assay (Progema, Cat# J8021) according to manufacture protocols. HRMEC cells were treated with *si-SLC38A5* or si-Ctrl at passage 6. Forty-eight hours after siRNA transfection, culture medium of each well was replaced with fresh culture medium at equal volume. 72-hour post siRNA treatment, both culture medium and cells were harvested for the glutamine/glutamate-Glo assay. Amounts of glutamine/glutamate in samples were determined from luminescence readings by comparison to a standard titration curve.

### SLC38A5 promoter cloning and dual-luciferase reporter assay

Cloning and reporter assays were performed based on previous protocols (55). Putative TCF-binding motif (A/TA/TCAAAG) was identified in three mouse *Slc38a5* promoter regions, which were amplified by PCR using the following primers: *Slc38a5_P1,* F: 5’-TATCGCTAGCCCAGCAGGGTGTATTTATG-3’ and R: 5’-TATCCTCGAGGGAGCGCTTTCAATCCTCAG-3’; *Slc38a5_P2,* F: 5’-TATCGCTAGC TCTCAAACTGTATCATGGAG-3’ and R: 5’-TATCCTCGAGCTACTTGCTGAAGACGTTG-3’; *Slc38a5_P3,* F: 5’-TATCGCTAGCAGGTCCTCTGAAGTATTGATC-3’ and R: 5’-TATCCTCGAGAGGAGAGTTCAAGTGTAGGT-3’. PCR products were purified by gel extraction, cloned into the pGL3 promoter luciferase vector (Promega, Madison, WI; E1751), and verified by Eton Bioscience (Boston, MA) with 100% matching sequence matching to the promoter region of *Slc38a5*. Both the promoter plasmids and a stabilized active form of β-catenin plasmid were transfected into HEK293T cells. After 48 hours, luciferase activity was measured with a dual-luciferase reporter assay kit (Promega; E1910), and the relative luciferase activity was determined by normalizing the firefly luciferase activity to the respective *Renilla* luciferase activity.

### RNA isolation and real-time RT-PCR

Total RNA was isolated from mouse retinas or from HRMEC culture with RNeasy Kit (Qiagen) based on manufacturer protocols. For LCM isolated vessels, RNA was isolated with the RNeasy Micro Kit (Qiagen, Cat# 74004). Synthesis of cDNA and q-PCR were performed using established standard protocols.

Primers for mouse real-time RT-PCR include: *Slc38a5:* F:5’-GACCTTTGGATACCTCACCTTC-3’ and R: 5’-CCAGACGCACACAAAGGATA-3’; 18s_F: 5’-CACGGACAGGATTGACAGATT-3’ and R: 5 ‘ - GCCAGAGTCTCGTTCGTTATC-3 ‘.

Human primers include: *SLC38A5:* F:5’-GAAGGGAAACCTCCTCATCATC-3’ and R: 5’-CAGGTAGCCCAAGTGTTTCA-3’; *18S_F:* 5’-GCCTCGAAAGAGTCCTGTATTG-3’ and R: 5’-TGAAGAGGGAGCCTGAGAAA-3’; *FLT1*(VEGFR1): F: 5’-CCGGCTCTCTATGAAAGTGAAG-3’ and R: 5’-CGAGTAGCCACGAGTCAAATAG-3’; KDR(VEGFR2): F: 5’-AGCAGGATGGCAAAGACTAC-3’ and R: 5’-TACTTCCTCCTCCTCCATACAG-3’; TEK(TIE2): F: 5’-TTTGCCCTCCTGGGTTTATG-3’ and R: 5’-CTTGTCCACTGCACCTTTCT-3’; *FGFR1:* F: 5’-GAGGCTACAAGGTCCGTTATG-3’ and R: 5’-GATGCTGCCGTACTCATTCT-3’; *FGFR2:* F: 5’-GGATAACAACACGCCTCTCTT-3’ and R: 5’ - CTTGCCCAGTGTCAGCTTAT −3’; *FGFR3*_F: 5’-CGAGGACAACGTGATGAAGA-3’ and R: 5’-TGTAGACTCGGTCAAACAAGG-3’; *IGF1R:* F:5’-CATGGTGGAGAACGACCATATC-3’ and R: 5’-GAGGAGTTCGATGCTGAAAGAA-3’; *IGF2R:* F: 5’ - CAGCGGATGAGCGTCATAAA-3’ and R: 5’-CGTGTCCCATGTGAAGAAGTAG-3’; *MAPK3(ERK1):* F: 5’-GCTGAACTCCAAGGGCTATAC-3’ and R: 5’-GTTGAGCTGATCCAGGTAGTG-3’; *MAPK1* (ERK2): F: 5’ - GGTACAGGGCTCCAGAAATTAT-3’ and R: 5’-TGGAAAGATGGGCCTGTTAG-3’; *MTOR:* F: 5’-GGGACTACAGGGAGAAGAAGAA-3’ and R: 5’ - GCATCAGAGTCAAGTGGTCATAG-3’.

### Western blot

Western blot was performed based on standard protocols (55). Retinal or HRMEC samples were extracted and sonicated in radioimmunoprecipitation assay lysis buffer (Thermo Fisher Scientific; 89901) with protease and phosphatase inhibitors (Sigma-Aldrich; P8465, P2850). Protein concentration was determined with a BCA protein assay kit. Equal amounts of protein were loaded in NuPAGE bis-tris protein gels (Thermo Fisher Scientific) and electroblotted to a polyvinylidene difluoride (PVDF) membrane. After blocking with 5% nonfat milk for 1 hour, membranes were incubated with a primary monoclonal antibody overnight at 4°C, followed by washing and incubation with secondary antibodies with horseradish peroxidase-conjugation (Amersham) for 1 hour at room temperature. Chemiluminescence was generated by incubation with enhanced chemiluminescence reagent (Thermo Fisher Scientific; 34075) and signal detected using an imaging system (17001401, Bio-Rad, Hercules, CA). Densitometry was analyzed with ImageJ software. Primary antibodies: anti-SLC38A5 (Biorbyt, St. Louis, MO; orb317962,), anti-non-phosphorylated β-catenin (Cell Signaling Technology; 8814S), anti-β-catenin (Santa Cruz Biotechnology, Santa Cruz, CA; sc-7199), anti–glyceraldehyde-3-phosphate dehydrogenase (GAPDH) (Santa Cruz Biotechnology; sc-32233), anti-VEGFR2 (Cell Signaling Technology; 2479). HRP-linked secondary antibodies were from Sigma-Aldrich: anti-mouse antibody (NA9310); anti-rabbit antibody (SAB3700934).

### Statistical analysis

Data were analyzed using GraphPad Prism 6.01 (GraphPad Software, San Diego, CA). Results are presented as means ± SEM (standard error of the mean) from animal studies; means ± SD (standard deviation) for non-animal studies with at least three independent experiments. N numbers listed in figure legends represent biological replication. Experimental groups were allocated based on genotypes for mutant mice work, eye position (left vs. right) for siRNA work, and randomly for in vitro work. Quantification analysis of in vivo retinal vascular phenotype were done with the identity of the samples masked. Statistical differences between groups were analyzed using a one-way analysis of variance (ANOVA) statistical test with Dunnett’s multiple comparisons tests (more than 2 groups) or two-tailed unpaired *t* tests (2 groups); *P* < 0.05 was considered statistically significant.

## Results

### *Slc38a5* expression is enriched in retinal blood vessels and down-regulated in Wnt signaling deficient *Lrp5^-/-^* and *Ndp^y/-^* retinal vessels

We found that the mRNA levels of *Slc38a5* were consistently down-regulated in both *Lrp5^-/-^* and *Ndp^y/-^* retinas lacking Wnt signaling from postnatal day (P) 5 through P17, compared with their age-matched WT controls (**Fig. 1A&B**). To localize the cellular source of *Slc38a5,* blood vessels were isolated from retinal cross-sections by laser capture microdissection (LCM), followed by mRNA expression analysis using RT-qPCR. There was ~40-fold enrichment of *Slc38a5* mRNA level in retinal blood vessels compared with that in the whole retinas in WT mice (**Fig. 1C**). Moreover, *Slc38a5* mRNA levels were substantially down-regulated in both *Lrp5^-/-^* and *Ndp^y/-^* LCM-isolated retinal vessels compared with their respective WT control blood vessels (**Fig. 1D**). Protein levels of *Slc38a5* were also significantly decreased in both P17 *Lrp5^-/-^* and *Ndp^y/-^* retinas versus their respective WT control retinas, with more substantial reduction in *Ndp^y/-^* retinas by ~50% (**Fig. 1E&F**).

**Figure 1:**
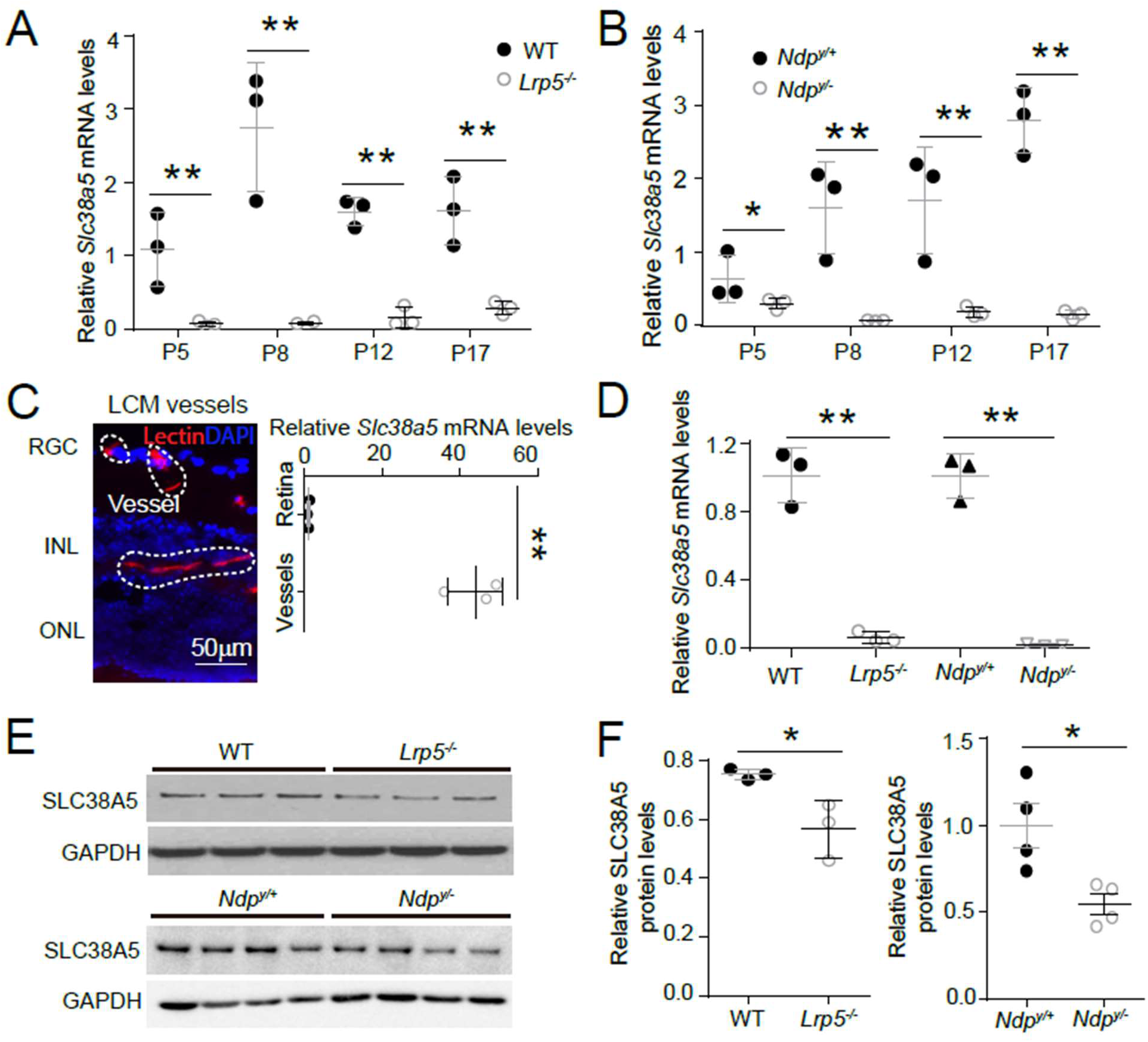
*Scl38a5* expression is enriched in retinal blood vessels and down-regulated in the retinas and retinal blood vessels of Wnt signaling deficient *Lrp5^-/-^* and *Ndp^y/-^* mice. **A-B.** mRNA levels of *Slc38a5* were measured by RT-PCR in *Lrp5^-/-^* (A) and *Ndp^y/-^* (B) retinas compared with their respective WT controls during development at postnatal day 5 (P5), P8, P12 and P17. **C.** Retinal blood vessels stained with isolectin B4 (red, outlined by the white dashed lines) were isolated by laser-captured microdissection (LCM) from retinal cross-sections. Cell nuclei were stained with DAPI (blue) for illustration purpose only. LCM retinal samples were stained with only isolectin B4 without DAPI. RGC: retinal ganglion cells. INL: inner nuclear layer. ONL: outer nuclear layer. *Slc38a5* mRNA levels in LCM isolated retinal blood vessels were compared with the whole retinal levels using RT-qPCR. Scale bars: 50 μm. **D.** *Slc38a5* mRNA levels in LCM-isolated *Lrp5^-/-^* and *Ndp^y/-^* retinal blood vessels were quantified with RT-qPCR and compared with their respective WT controls. **E-F:** Protein levels of SLC38A5 in P17 *Lrp5^-/-^* and *Ndp^y/-^* retinas and their WT controls were quantified with Western blot (E), and normalized by GAPDH levels (F). Data are expressed as mean ± SEM. n = 3-4 per group. *p ≤ 0.05, **p ≤ 0.01.

*Slc38a5* expression in retinal cells was further analyzed in single cell transcriptome datasets of P14 C57B6 mouse retinas (56), where *Slc38a5* is found mainly expressed in vascular endothelium, similar as endothelium marker *Pecam1* (**Fig. S1A**). In a human retinal single cell dataset - Cell atlas of the human fovea and peripheral retina (57), *SLC38A5* is also mainly expressed in vascular endothelium, just like *PECAM1* (**Fig. S1B**).

Together these data demonstrate significant down-regulation of *Slc38a5* mRNA and protein levels in *Lrp5^-/-^* and *Ndp^y/-^* retinas, which is consistent with previous gene array findings from our and others’ studies (38, 40). These findings also strongly support vascular endothelium specificity of *Slc35a5* and its potential role in angiogenesis.

### Wnt signaling directly regulates *Slc38a5* transcription in vascular endothelium

To assess whether modulation of Wnt signaling directly regulates *Slc38a5* expression in retinal vascular endothelium, human retinal microvascular endothelial cells (HRMECs) were cultured and treated with recombinant Norrin, a Wnt ligand, in the presence or absence of a Wnt inhibitor XAV939. XAV939 is a small molecule inhibitor of the Wnt signaling pathway, and it works through binding to tankyrase and stabilizing the Axin proteins, thus increasing degradation of β-catenin and blocking Wnt signaling (58). We found that *Slc38a5* mRNA and protein levels were substantially induced by Norrin by ~1.6 fold (**Fig. 2A&B**), whereas subsequent treatment with XAV939 reversed the Norrin-induced SLC38A5 upregulation in HRMECs (**Fig. 2A&B**). Moreover, Wnt3a conditioned medium (Wnt3a-CM) induced an even more potent upregulation of SLC38A5 by ~4-fold in mRNA levels and ~10 fold in protein levels (**Fig. 2A&C**), which were also reversed by XAV939 treatment (**Fig. 2 A&C**). Activation or inhibition of Wnt signaling by Wnt modulators (Norrin, Wnt3a-CM and XAV939) were confirmed in Western blot by the levels of the active β-catenin (non-phosphorylated-β-catenin), the canonical Wnt effector (**Fig. 2B&C**).

**Figure 2:**
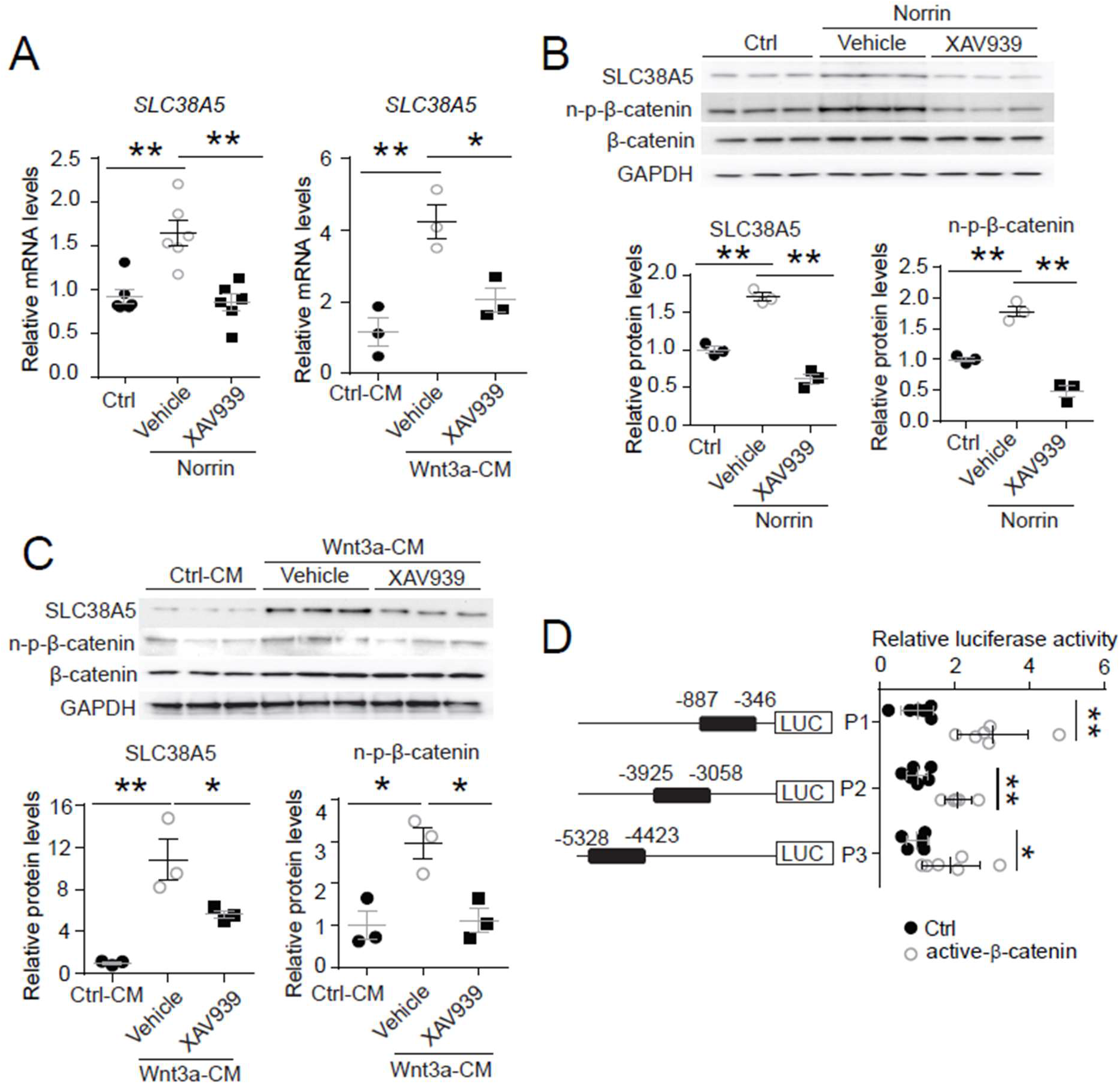
*Slc38a5* is a direct target gene of Wnt signaling in the vascular endothelium. **A:** *Slc38a5* mRNA levels were increased in human retinal microvascular endothelial cells (HRMECs) treated with Wnt ligands, recombinant Norrin and Wnt3a-conditioned medium (Wnt3a-CM), compared with their respective vehicle controls (Ctrl, and Ctrl-CM), and suppressed by a Wnt inhibitor XAV939. **B-C:** Protein levels of SLC38A5 in HRMECs were up-regulated by Wnt ligands Norrin (B) and Wnt3a-CM (C), and down-regulated by XAV939. Protein levels of SLC38A5 were quantified by Western blotting and normalized by GAPDH levels. n-p-β-catenin: non-phosphorylated β-catenin. **D:** Three promoter regions upstream of *Slc38a5* gene containing potential Wnt-responsive TCF-binding motifs (TTCAAAG) were identified based on sequence analysis. Three putative TCF binding regions: P1 (−887 bp to −346 bp), P2 (−3925 bp to −3058 bp) and P3 (−5328 bp to −4423 bp) were cloned and ligated separately with a luciferase reporter, and co-transfected with an active β-catenin plasmid in HEK 293T cells, followed by measurement of luciferase activity. Data are expressed as mean ± SEM. n = 3-6 per group. *p ≤ 0.05, **p ≤ 0.01.

To determine whether Wnt signaling directly regulates *SLC38A5* expression at the transcription level, dual luciferase reporter assays were constructed in HEK293T cells. Canonical Wnt signaling mediates transcription of its target genes through recognizing β-catenin-responsive TCF-binding motifs (A/TA/TCAAAG) on their regulatory regions (59). Three *SLC38A5* promoter regions containing putative TCF-binding motifs were identified, cloned and ligated with a luciferase reporter. All three luciferase reporter-containing promotor regions: P1, P2, and P3, showed significant increase in luciferase activity when co-transfected with active β-catenin, with both P1 and P2 showing ~3 fold increase and P3 showing ~2-fold increase (**Fig. 2D**). Together, these results suggest that SLC38A5 transcription is directly regulated by Wnt signaling via β-catenin binding to potentially multiple TCF-binding sites on its promoter regions.

### Genetic deficiency of *Slc38a5* impairs developmental retinal angiogenesis

To evaluate the role of *Slc38a5* in retinal vessel development, we injected intravitreal siRNA targeting *Slc38a5* (si-*Slc38a5*) and control negative siRNA (si-Ctrl) to developing C57BL/6J mouse eyes and analyzed the impact on vascular development. Treatment with siRNA resulted in substantial reduction of *Slc38a5* mRNA levels by ~70% (n=3/group, p≤ 0.01, **Fig. 3A**) and protein levels by ~70% (n=3/group, p≤ 0.05, **Fig. 3B**) compared with contralateral si-Ctrl injected eyes at 3 days post injection. Moreover, si- *Slc38a5* treated retinas showed significant (~20%) delay of superficial vascular coverage in the retinas at P7, 3 days after injection at P4, compared with si-Ctrl injection (n=9/group, p≤ 0.05, **Fig. 3C**). In addition, the development of deep retinal vascular layer was also substantially impaired with almost 40% reduction in deep layer vascular area at P10, 3 days after siRNA were injected at P7 (n=11/group, p≤ 0.01, **Fig. 3D**). These data suggest that knockdown of *Slc38a5* expression by siRNA delivery impedes retinal vascular development.

**Figure 3.**
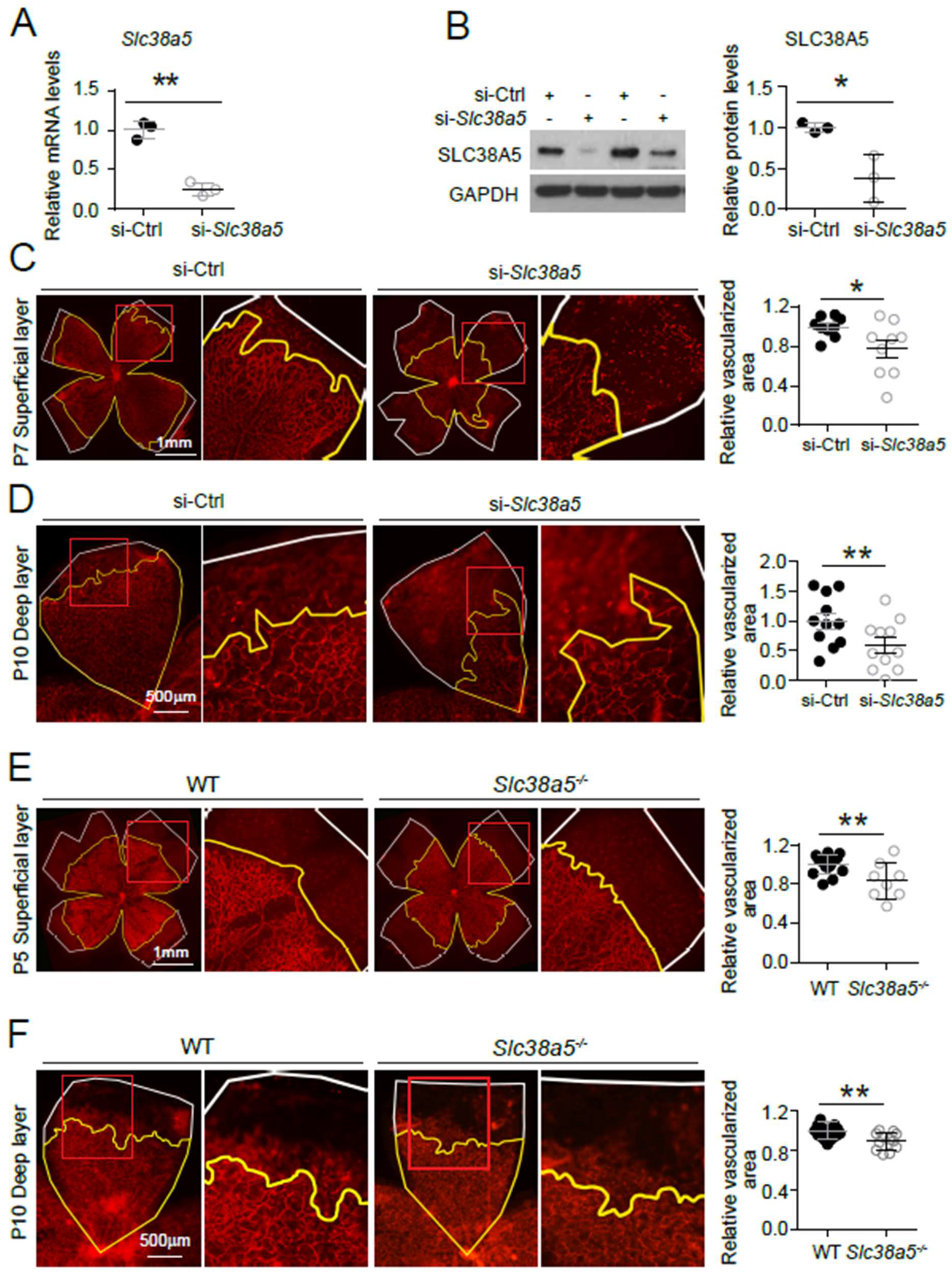
Genetic deficiency of *Slc38a5* impairs developmental retinal angiogenesis in vivo. **A-D:** siRNA targeting *Slc38a5* (si-*Slc38a5*) was intravitreally injected in C57BL/6J mice (WT), and the same volume of negative control siRNA (si-Ctrl) was injected into the contralateral eyes. Mice were sacrificed 3 days after injection and retinas were isolated to detect expression level or to quantify vascular growth. mRNA (A) and protein (B) levels of SLC38A5 confirms successful knockdown. Each lane represents 1 retina. Retinal vascular coverage of superficial layer at P7 (C) and deep layer at P10 (D) were analyzed 3 days after intravitreal injection of si- *Slc38a5* and compared with their respective controls. Retinas were dissected, stained with isolectin B4 (red), and then flat-mounted to visualize the vasculature. Percentages of vascularized area were quantified in superficial (C, n=9/group) or deep (D, n=11/group) vascular layer. **E&F:** Retinal blood vessel development in *Slc38a5^-/-^* and WT littermate control mice from the same colony was imaged and quantified at P5 (E, n=8-11/group) and P10 (F, n=12-13/group), with staining of isolectin B4 (red) to visualize the vasculature. In panels C-F, yellow lines outline retinal vascular areas and white lines indicate total retinal areas. Red boxes indicate location of enlarged insets as shown on the right. Each dot represents one retina. Data are expressed as individual value and mean ± SEM. *p ≤ 0.05, **p ≤ 0.01.

The role of *Slc38a5* in retinal vessel development was further evaluated in mutant mice with genetic deficiency of *Slc38a5.* Mice with targeted disruption of the *Slc38a5* gene are grossly normal yet have decreased pancreatic alpha cell hyperplasia induced by glucagon receptor inhibition (35). Retinal blood vessel development was substantially delayed in *Slc38a5^-/-^* retinas for superficial vascular layer at P5 (n=8-11/group, p≤ 0.01, **Fig. 3E**) and deep vascular layer at P10 (n=12-13/group, p≤ 0.01, **Fig. 3F**). The delayed vascular development, however, is resolved in adult *Slc38a5^-/-^* mice when the retinal vasculature are largely normal in both superficial and deep layers (**Fig. S2A&C**). These findings suggest that genetic knockout of *Slc38a5* results in delayed vascular development in the retinas and highlight an important role of *Slc38a5* in normal retinal vessel development.

### SLC38A5 is enriched in pathological neovessels and its genetic deficiency dampens pathological angiogenesis in OIR

To evaluate the role of *Slc38a5* in pathological retinal angiogenesis, we used a well-established oxygen-induced retinopathy (OIR) model (45), mimicking the hypoxia-induced proliferative phase as seen in retinopathy of prematurity and diabetic retinopathy. In this model, neonatal mice and their nursing mother were exposed to 75±2% oxygen from P7 to P12 to induce vaso-obliteration, followed by return to room air, when relative hypoxia induces vaso-proliferation with maximal neovessel formation observable at P17. We found that *Slc38a5* mRNA expression was significantly suppressed during the vaso-obliteration phase between P8 and P12, and up-regulated during the vaso-proliferative phase of OIR between P14 and P17, compared with age-matched room air control mice (**Fig. 4A**). Enrichment of *Slc38a5* in pathological neovessels was also quantitatively measured in OIR neovessels vs. normal vessels isolated using laser capture micro-dissection. In pathological neovessels from P17 OIR retinas, *Slc38a5* mRNA levels were enriched at ~2.5-fold, compared with the levels in age-matched normoxic vessels (**Fig. 4B**). Moreover, SLC38A5 protein levels were also increased at P17 OIR whole retinas by ~2 fold compared with age-matched normoxic retinas in Western blot (**Fig. 4C**). These data indicate that *Slc38a5* is up-regulated in pathological retinal neovessels and suggestive of its angiogenic role in retinopathy.

**Figure 4.**
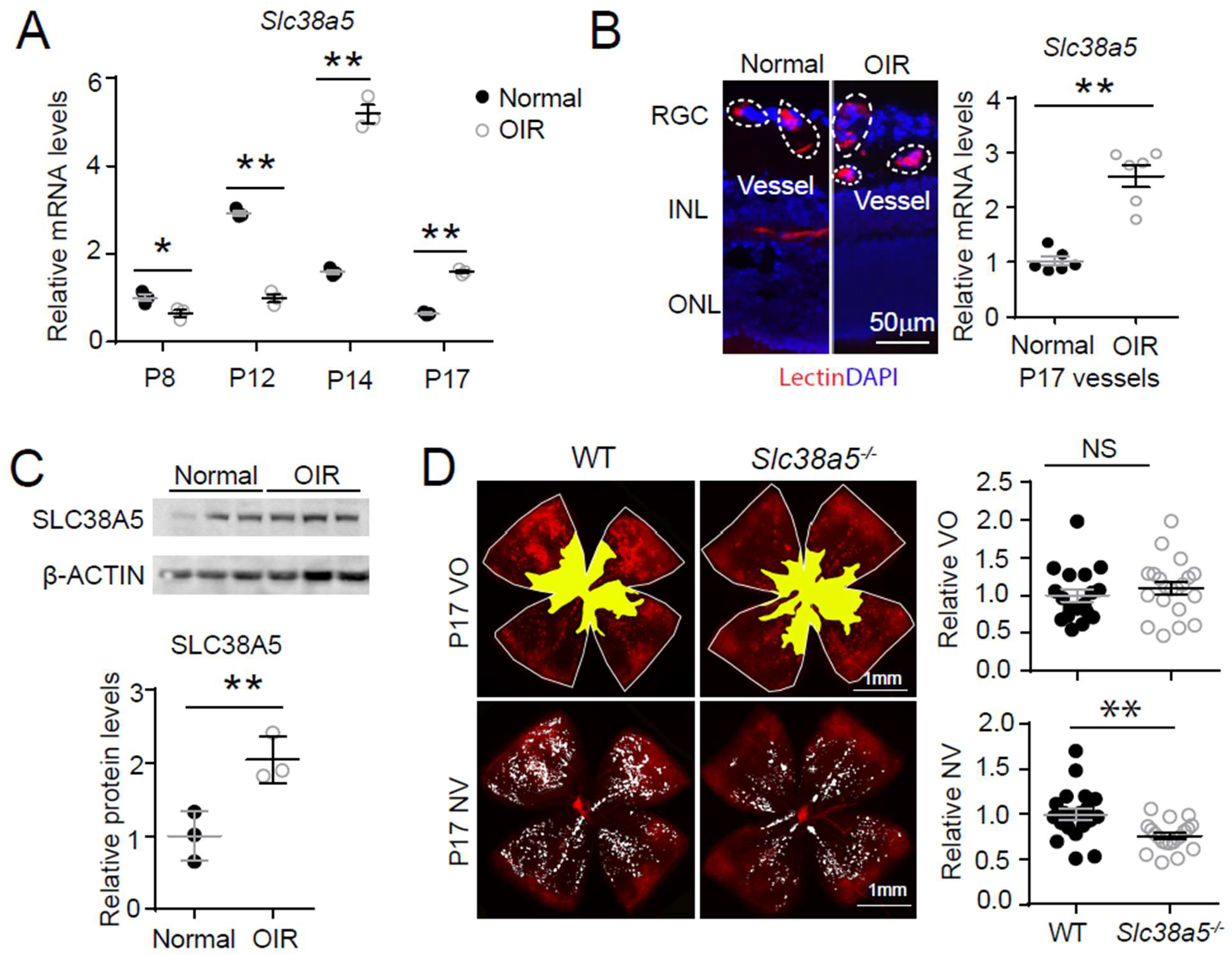
*Slc38a5* is enriched in OIR pathological neovessels and its deficiency suppresses pathological angiogenesis in OIR. P **A:** *Slc38a5* mRNA expression was measured by RT-qPCR at P8, P12, P14 and P17 in C57BL/6J OIR retinas compared with age-matched normoxic control mice. *Slc38a5* mRNA levels were decreased during hyperoxia stage (P8 and P12) and increased in hypoxia stage (P14 and P17). **B:** *Slc38a5* mRNA expression was analyzed using RT-qPCR in laser capture micro-dissected (LCM) pathological neovessels from P17 unfixed C57BL/6J OIR retinas compared with normal vessels isolated from P17 normoxic retinas. Images on the left are representative retinal cross-sections from normal and OIR retinas stained with isolectin B4 (red) and DAPI (blue), with dotted lines highlighting micro-dissected retinal vessels. GCL: ganglion cell layer, INL: inner nuclear layer, ONL: outer nuclear layer. **C:** Protein levels of SLC38A5 were increased in C57BL/6J OIR retinas at P17 compared with normoxic controls using Western blot and quantified with densitometry. **D:** *Slc38a5^-/-^* exposed to OIR had decreased levels of pathological NV compared with WT OIR controls bred in the same colony at P17. There was no significant difference in vaso-obliteration between the two groups. Scale bar: 50 μm (**B**), 1 mm (**D**). Each dot represents one retina. Data are expressed as mean ± SEM. n = 3-6 per group (**A-C**), n = 20 per group (**D**). *p ≤ 0.05; **p ≤ 0.01; n.s.: not significant.

To further investigate the role of OIR-induced *Slc38a5* up-regulation in pathological neovessels, we subjected *Slc38a5^-/-^* mice to OIR. *Slc38a5^-/-^* mice showed significantly decreased levels of pathological neovascularization at P17 by ~25% as compared with WT littermate controls (n=20/group, p≤ 0.01, **Figure 4D**), whereas the vaso-obliterated retinal areas were comparable at P17. These data suggest that loss of *Slc38a5* leads to significantly decreased levels of pathological neovascularization in OIR.

### Modulation of SLC38A5 regulates EC angiogenic function *in vitro*

The function of SLC38A5 in mediating EC angiogenesis was assessed in HRMEC culture using siRNA to knockdown *SLC38A5*. Compared with negative control siRNA (si-Ctrl), *SLC38A5* siRNA (si- *SLC38A5*) effectively suppressed *SLC38A5* mRNA expression by more than 80% (p≤ 0.01, **Fig. 5A**), and protein level by ~50% (p≤ 0.05, **Fig. 5B**), confirming successful inhibition. EC viability and/or proliferation were analyzed with MTT cellular metabolic activity assay (measuring NAD(P)H-dependent oxidoreductase activities) at 1-3 days after siRNA transfection. HRMEC metabolic activity was significantly decreased at 48 and 72 hours after transfected with si- *SLC38A5* (~30% reduction at 72 hours), compared with si-Ctrl-treated HRMEC (p≤ 0.01, **Fig. 5C**), indicative of decreased cell viability and/or proliferation. In addition, si-*SLC38A5* substantially suppressed HRMEC migration ((p≤ 0.05, **Fig. 5D**) and tubular formation, resulting in ~35% reduction in total vessel length (**Fig. 5E**). Together these results suggest that knockdown of amino acid transporter SLC38A5 potent suppresses EC angiogenic function.

**Figure 5.**
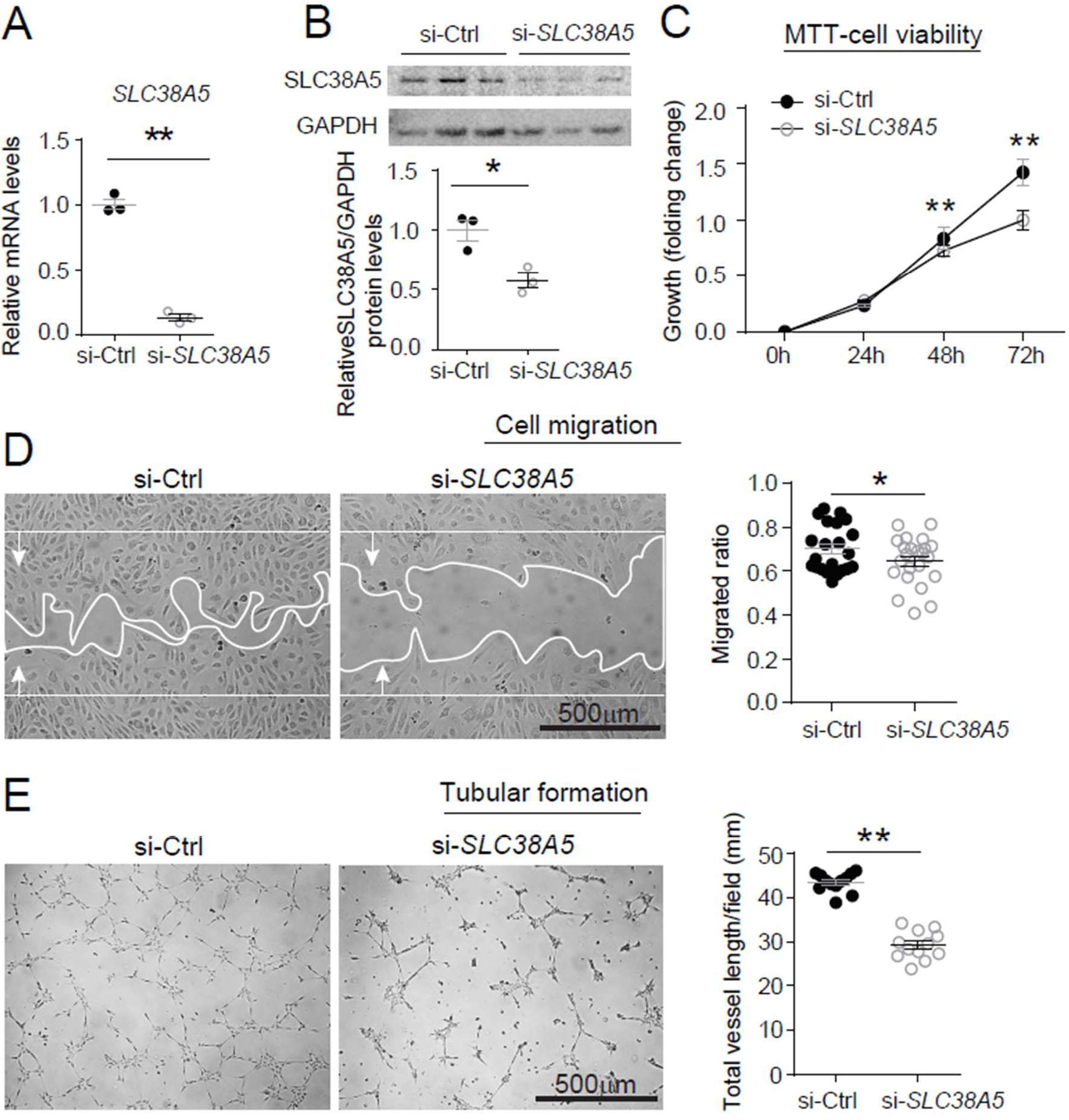
Inhibition of *SLC38A5* dampens endothelial cell viability, migration and tubular formation in vitro. HRMECs were transfected with siRNA targeting *SLC38A5* (si-*SLC38A5*) or control siRNA (si-Ctrl). **A-B:** mRNA (A) and protein (B) levels of *SLC38A5* confirm successful knock down by si-*SLC38A5*. **C:** HRMEC cell viability was measured with MTT assay. Cell growth rate was calculated as fold change normalized to the values at 0 hour. **D:** HRMECs were grown to confluence and treated with si- *SLC38A5* or si-Ctrl for 48 h. Cells were then treated with mitomycin to stop cell proliferation. A scratch was performed to the cells to generate a wound. Migrated areas (new cell growth areas normalized by original wound areas) of HRMECs were measured after 16 h. **E.** Tubular formation assay was conducted by collecting cells after 48 hours of si- *SLC38A5* transfection, and seeding cells onto Matrigel-coated wells to grow for additional 9 hours. Representative images show formation of EC tubular network and total vessel length per field was analyzed by Image J. Scale bar: 500μm (**D&E**). Data are shown as mean ± SEM; n=3-6/group. *p ≤ 0.05; **p ≤ 0.01.

### SLC38A5 inhibition decreases EC glutamine uptake

As an AA transporter, one of the main AA that SLC38A5 transports is glutamine. Here, the impact of SLC38A5 on EC uptake of glutamine was measured with a glutamine/glutamate uptake bioluminescent assay in HRMECs. Suppression of SLC38A5 with si-*SLC38A5* resulted in ~25% decrease in the amount of intracellular glutamine measured by bioluminescence and then titrated to a standard concentration curve, compared with si-Ctrl treatment (p≤ 0.01, **Fig. 6A**). Conversely, measurement of corresponding cell culture medium collected from si-*SLC38A5*-treated HRMECs also shows a reciprocal ~25% higher levels of glutamine content than si-Ctrl-treated cells ((p≤ 0.01, **Fig. 6B**), suggesting effective blockade of glutamine transport into HRMECs.

**Figure 6.**
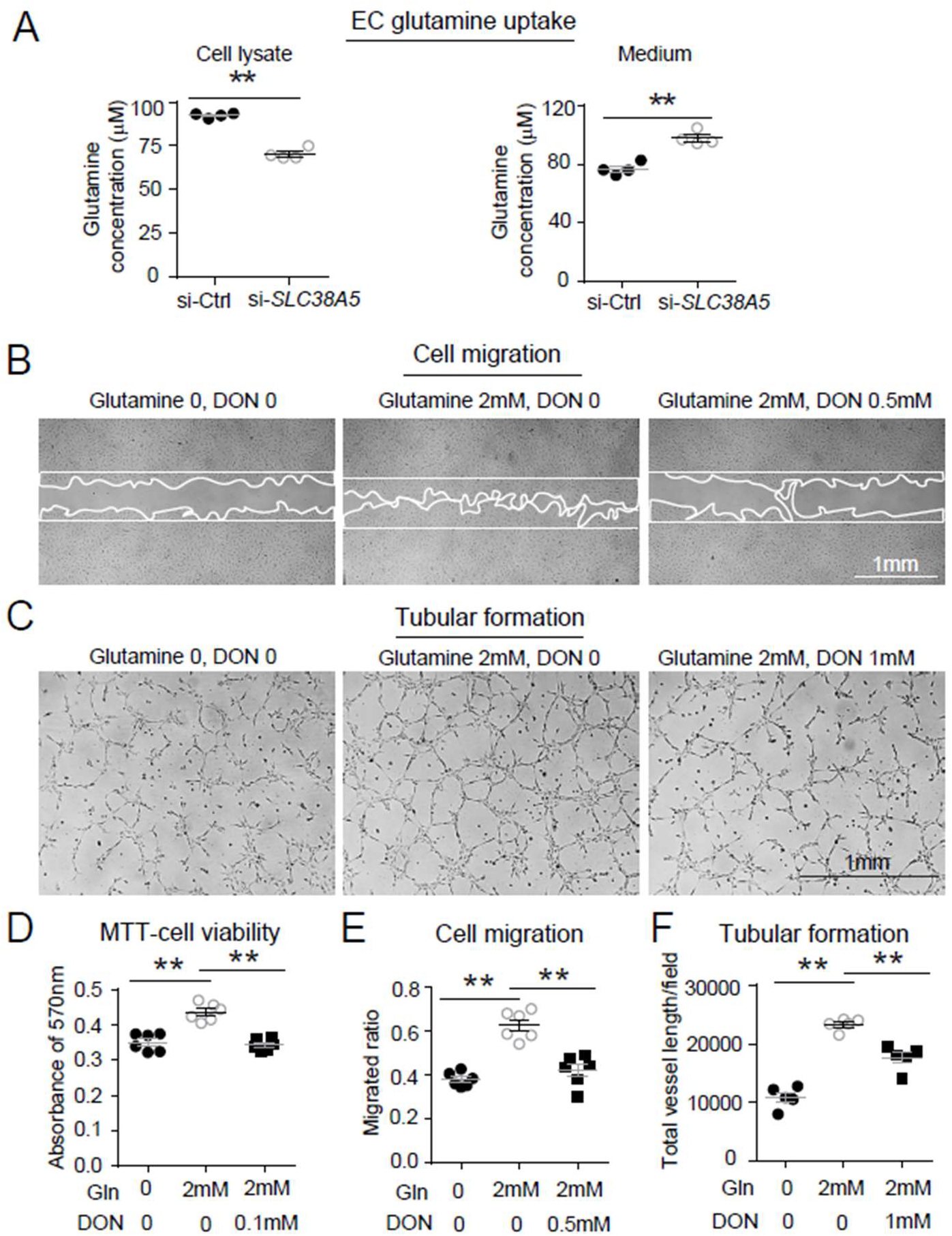
SLC38A5 facilitates EC uptake of glutamine, which is essential for EC viability, migration, and tubular formation. **A:** SLC38A5 knockdown with si-*SLC38A5* suppressed glutamine uptake by HRMECs, with decreased glutamine levels in HRMEC cell lysates and increased culture medium levels, measured with a glutamine/glutamate-Glo bioluminescent assay. Levels of glutamine/glutamate in HRMECs and culture medium samples were determined from bioluminescence readings by comparison to a standard titration curve. **B & E:** HRMECs were grown to confluence and a scratch was applied to generate a wound. Mitomycin was used to stop cell proliferation. A glutamine antagonist, 6-diazo-5-oxo-norleucine (DON), was used to broadly inhibit glutamine uptake. 16 hours were given to the cells to migrate. Representative images are shown in (B) and the quantification of migrated areas are shown in (E). **C, F:** HRMECs treated were seeded onto Matrigel for 9 hours and treated with glutamine and DON for tubular formation. Representative images are shown in (C) and the quantification of total vessel length per field are shown in (F). **D:** HRMEC cell viability was measured at 24 hours by MTT assay and normalized to the levels at 0 hour to quantify the cell growth rate. Scale bars: 1 mm (**B&C**). Data are expressed as means ± SEM. n = 4-6 per group. *p ≤ 0.05; **p ≤ 0.01.

The effect of glutamine on HRMEC angiogenesis was then further examined with angiogenic assays. Treatment with glutamine enhanced HRMEC proliferation in MTT assays, which was blocked by a glutamine antagonist 6-diazo-5-oxo-L-norleucine (DON) (**Fig. 6D**). DON has structural similarity with glutamine and hence can lead to competitive binding of glutamine binding proteins and their irreversible inhibition by forming a covalent adduct (60). Moreover, glutamine treatment substantially increased HRMEC migration by ~50% (**Fig. 6 B&E**) and tubular formation by more than 2-fold (**Fig. 6C&F**), both of which were reversed by DON treatment. These findings further suggest that loss of SLC38A5 may limit glutamine uptake and reduce its bioavailability in vascular EC, thereby leading to decreased EC angiogenesis.

### Suppression of SLC38A5 alters key pro-angiogenic factor receptor and signaling in ECs

To understand the factors mediating the effect of SLC38A5 on EC angiogenesis, we evaluated expression of multiple angiogenic factor receptors in response to SLC38A5 knockdown. Suppression of SLC38A5 led to substantially altered mRNA expression of VEGF receptors (VEGFR1 and VEGFR2), Tie2, FGF receptors (FGFR1-3), IGF1R and IGF2R (**Fig. 7A**). In addition, there was substantial suppression of expression of ERK1&2 and mTOR (**Fig. 7A**), key mediators downstream of VEGFR2 signaling that control EC growth. As VEGF-A is one of main angiogenic growth factors, we measured the protein levels of VEGF receptors including VEGFR2, the major angiogenic receptor for VEGF-A, and VEGFR1, often acting as a VEGF decoy trap to modulate VEGFR2 function (61). Protein levels of VEGFR1 were substantially up-regulated, whereas VEGFR2 levels were substantially suppressed after SLC38A5 knockdown in HRMECs (**Fig. 7B**). These findings suggest that suppression of SLC38A5 dampens EC glutamine uptake, and subsequently leads to altered expression of growth factor receptors such as VEGFR1&2 to restrict EC angiogenesis.

**Figure 7.**
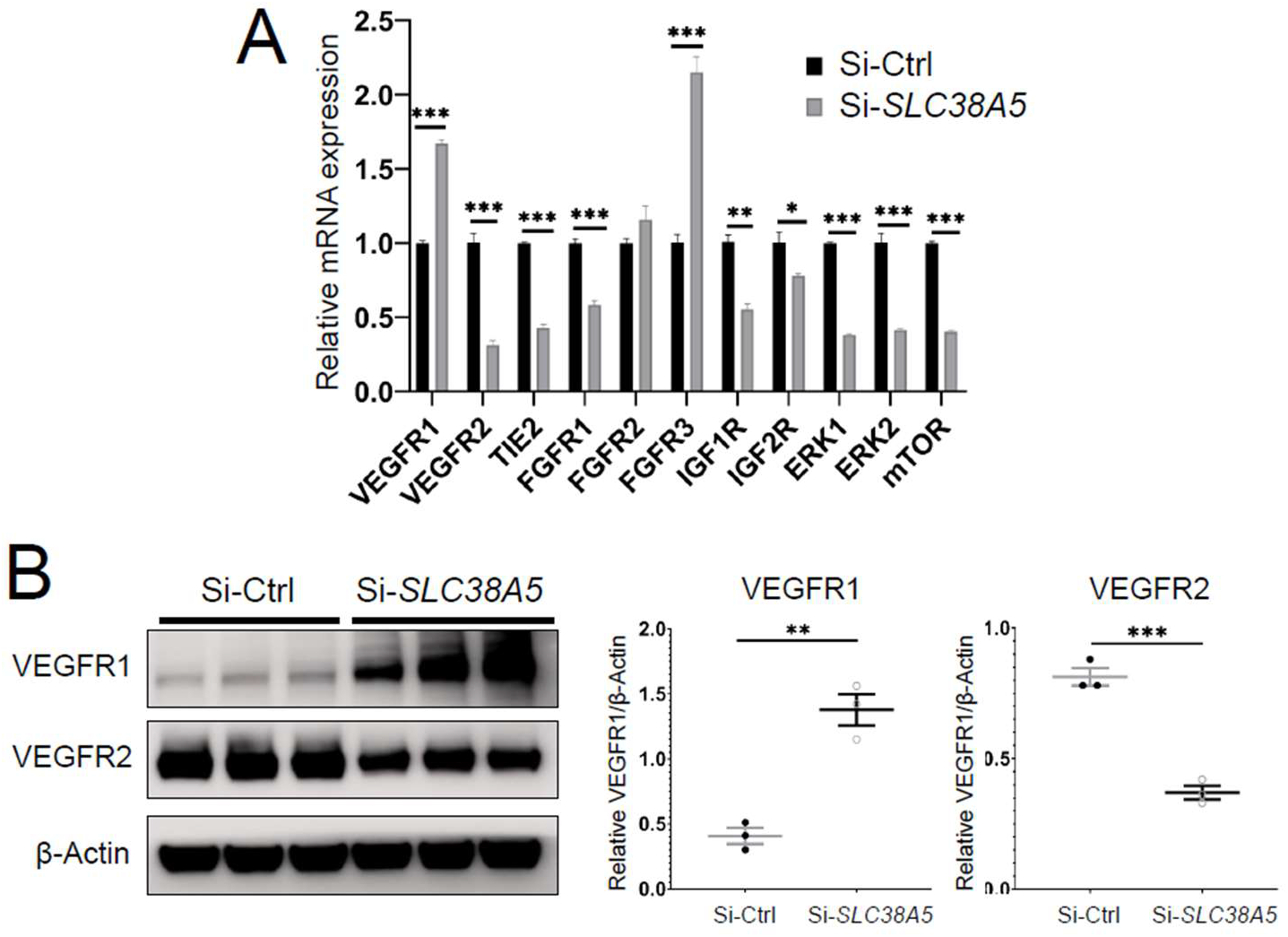
Suppression of Slc38a5 modulates growth factor receptors including VEGFR1 and VEGFR2. HRMECs were transfected with siRNA targeting *SLC38A5* (si-*SLC38A5*) or control siRNA (si-Ctrl) for 72 hours, and collected for RT-qPCR or Western blots. **A:** mRNA levels of growth factor receptors and signaling molecules were normalized by expression of 18S (n=3-6/group). **B.** Western Blots show protein levels of VEGFR1 and VEGFR2 with Si-SLC38A5 or si-Ctrl treatment. Data are shown as mean ± SEM; n=3/group. **p ≤ 0.01; ***p ≤ 0.001.

## Discussion

This study establishes that amino acid transporter SLC38A5 is a new pro-angiogenic regulator in the retinal vascular endothelium both during development and in retinopathy. SLC38A5 transcription is regulated by Wnt signaling, a fundamentally important pathway in retinal angiogenesis (41, 62). We demonstrate that SLC38A5 deficient retinas have delayed developmental angiogenesis and dampened pathological neovascularization in an oxygen-induced retinopathy. Based on these findings we suggest that SLC38A5 regulates retinal angiogenesis through modulating vascular EC uptake of amino acids such as glutamine, and thereby influencing angiogenic receptor signaling including VEGFR2 to impact EC growth and function (**Fig. 8**).

**Figure 8.**
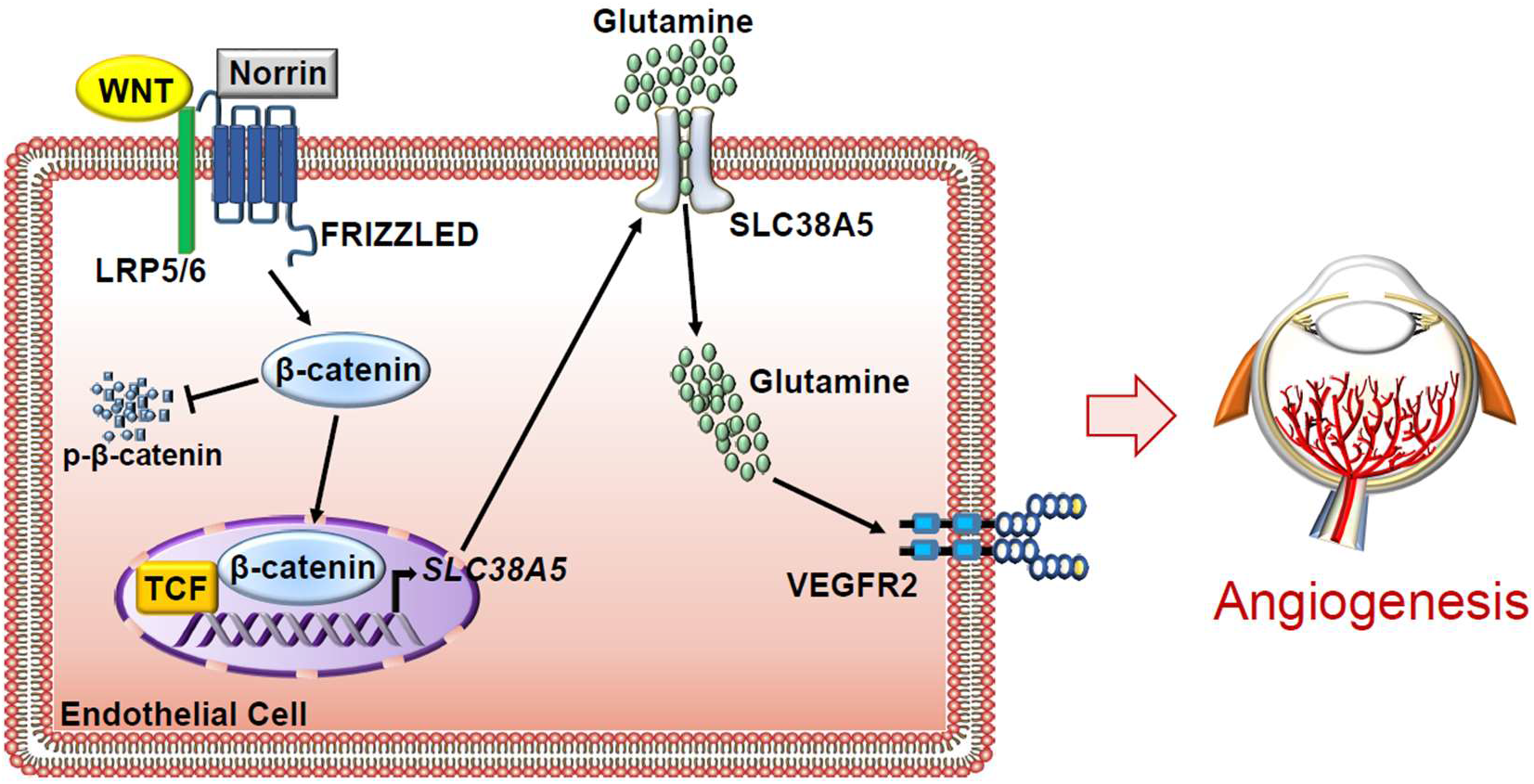
Schematic illustration of a pro-angiogenic role of amino acid transporter SLC38A5 in retinal angiogenesis. In vascular ECs, Wnt ligands (Wnts and Norrin) activate Wnt/β-catenin signaling, which directly controls the transcription of EC-enriched SLC38A5 by binding to a TCF binding site on SLC38A5 promoter. Endothelial SLC38A5 facilitates EC uptake of AAs such as glutamine as energy fuel and source of protein synthesis. Altered glutamine and nutrient availability in EC subsequent affects VEGFR2 levels and signaling, and thus retinal angiogenesis. In retinopathy, expression of both Wnt receptors and endothelial SLC38A5 are enriched in pathological neovessels, promoting glutamine availability and thereby contributing to VEGFR2 signaling and formation of pathologic retinal neovascularization. Inhibition of SLC38A5 may suppress pathologic neovessels and alleviate pathologic neovascularization in retinopathy.

Previously studies identified SLC38A5 localization in brain glial cells (32) and in the retinal Müller glia and retinal ganglion cells (36, 37), yet studies on its function in the brain and eyes are scarce, other than work on its intrinsic role as a glutamine transporter to provide precursor for the neurotransmitter glutamate (63, 64). Elsewhere in the body, identification of SLC38A5 as a marker of pancreatic alpha cell precursor expanded its role in regulating alpha cell proliferation and hyperplasia through nutrient sensing (33–35). In the intestine, SLC38A5 was found in crypt cells and may participate in chronic intestinal inflammation (65, 66). Here, our findings with laser capture microdissection and single cell transcriptome analysis identified specific enrichment of *Slc38a5* in retinal vascular endothelium, thus uncovering its new role as an angiogenic regulator and a new marker of sprouting neovessels.

Upregulation of *Slc38a5* in retinal neovessels is potentially regulated by the Wnt/β-catenin signaling pathway, as indicated by our findings in *Lrp5^-/-^* and *Ndp^y/-^* mice and in EC cell culture. We showed that *Slc38a5* is a new direct target gene of Wnt/β-catenin signaling, and that it is deficient in both *Lrp5^-/-^* and *Ndp^y/-^* retinas and blood vessels. Our prior work found that Wnt signaling is enriched in developing and pathological neovessels in OIR (67), further supporting the notion that the Wnt pathway induces *Slc38a5* transcription in neovessels to promote angiogenesis. Delayed development of *Slc38a5^-/-^* retinal vessels, however, are resolved by adult age with normal vasculature and intact deep layer of vessels (**Fig. S2**), indicating that *Slc38a5^-/-^* retinas do not reproduce FEVR-like symptoms as seen in *Lrp5^-/-^* and *Ndp^y/-^* mice. This suggests that down regulation of *Slc38a5* may only partially explain the effects of Wnt signaling on angiogenesis, and other Wnt- and β-catenin-mediated factors are still at work to drive defective angiogenesis in *Lrp5^-/-^* and *Ndp^y/-^* retinas, including for example, Sox family proteins (68) and integrin-linked kinase (ILK) (69). In addition to regulating retinal vessel growth, Wnt/β-catenin signaling is critical to maintain blood vessel barrier and prevent vascular leakage in the brain and eyes (41, 62, 70, 71). *Slc38a5^-/-^* eyes exhibit normal blood-retinal barrier with no detectable vascular leakage in fundus fluorescent imaging (**Fig. S2B**), suggesting that SLC38A5 is dispensable in mediating the effects of Wnt signaling on the retinal vascular barrier largely through both tight junctions such as claudin 5 (38, 67), and regulation of EC transcytosis (55). Other than Wnt signaling, *Slc38a5* may potentially be responsive to oxygen sensing and regulated by hypoxia/HIF, as *Slc38a5* expression is suppressed in the phase I of OIR during oxygen exposure and upregulated in the second phase of relative hypoxia.

Our data showed that glutamine uptake in vascular endothelium is substantially down-regulated when SLC38A5 is suppressed, suggesting that SLC38A5 may regulate glutamine availability and thus control EC angiogenesis. Metabolic regulation of blood vessels has been increasingly recognized to play important roles in driving angiogenesis (72, 73). Specifically, amino acids including glutamine and asparagine can drive EC proliferation and vessel sprouting through regulation of TCA cycle and protein synthesis (20, 21), since either depletion of glutamine or inhibition of glutaminase 1 (an enzyme that converts glutamine to glutamate), or suppression of asparagine synthetase, impairs angiogenesis (20, 21). Moreover, glutamine synthetase (another enzyme responsible for glutamine formation) promotes angiogenesis through activating small GTPase RHOJ, independent of its role in glutamine synthesis (74). Our findings on SLC38A5 and its transport of glutamine in EC, together with the effect of glutamine on EC angiogenic function, further strengthen the idea that glutamine is key to sprouting angiogenesis and suggest its bioavailability in EC is controlled in part by SLC38A5. Glutamine was reported previously to fuel EC proliferation but not migration in HUVEC *in vitro* (21), yet in our study we found glutamine promotes migration of HRMECs, which may reflect different cell-specific responses to glutamine in macrovascular HUVECs vs. microvascular HRMECs, as well as different migration assay methods used.

While glutamine is the most abundant AA in circulation, Müller glia also provide glutamine locally in the retina through uptake of and synthesis from excess extracellular glutamate as neurotransmitter (75). Thus SLC38A5 may help vascular ECs to uptake and utilize free glutamine from both systemic circulation and those released by Müller glia. A recent study found that hyperoxia promotes glutamine consumption and glutamine-fueled anaplerosis in Müller glia (76), suggesting a link between oxygen sensing and glutamine metabolism in the Müller glia. Whether SLC38A5 may interlink the oxygen-sensing and glutamine metabolism in Müller cells and ECs under other similar physiological or pathological conditions will await further studies.

In addition to glutamine, SLC38A5 may transport other AAs such as serine and glycine, which may further regulate angiogenesis in eye diseases. For example, serine deprivation in the absence of a synthesizing enzyme phosphoglycerate dehydrogenase (PHGDH) causes lethal vascular defects in mice (77), whereas serine-glycine metabolism is also required in angiogenesis and linked to endothelial dysfunction in response to oxidized phospholipids (78). In the mouse model of OIR, serine and one-carbon metabolism were found to mediate HIF response in retinopathy through liver-eye serine crosstalk (79). In macular telangiectasia type 2 (MacTel), a rare degenerative eye disease with abnormal intraretinal angiogenesis, defective serine biosynthesis and PHDGH haploinsufficiency were genetically linked with disease onset and associated with toxic ceramide accumulation in circulation and in retinal pigment epithelium cells (80, 81). Whether SLC38A5 may regulate serine transportation to influence retinal angiogenesis and retinal diseases will need additional investigation.

We showed here that decreased glutamine uptake in EC is directly associated with altered expression of growth factor receptors and particularly increased levels of VEGFR1 and decreased levels of VEGFR2. As a major angiogenic growth factor, VEGF-A exerts its effects on ECs mainly through its receptor VEGFR2 to promote angiogenesis – including EC proliferation, migration and cellular differentiation (82), whereas VEGFR1 often has negative angiogenic function and acts largely as a decoy receptor of VEGF to limit VEGFR2 function (61). Suppression of SLC38A5 may thus not only directly decrease VEGFR2 expression but also limit VEGFR2 function due to increased VEGFR1 levels, both of which can result in decreased retinal angiogenesis.

Findings presented here suggest that modulation of SLC38A5 and its associated AAs may have translational value in treating retinopathy. Premature infants often lack conditionally essential AAs such as glutamine or arginine (83, 84), due to their impaired endogenous synthesis. Providing much needed AAs early on might promote normalization of delayed vessel growth, and thereby preventing the neovascular phase of retinopathy. Previously, supplement of an arginine-glutamine (Arg-Glu) dipeptide was found to dampen pathological neovascularization in OIR by promoting normal vascular restoration (85, 86). Here our data suggest lack of SLC38A5, and likely subsequent impaired glutamine uptake, dampens developmental angiogenesis, in line with the pro-angiogenic role of AAs in the first phase of retinopathy. In the late proliferative phase of retinopathy, however, inhibiting of AA transporters like SLC38A5 or starving retinal vessels from AA nutrients may be beneficial in directly suppressing uncontrolled pathological neovascularization. Targeting of AAs and their transporters such as SLC38A5 may thus represent a new potential approach to treat retinopathy.

One limitation of the current study lies in the mutant mice used, which have systemic knockout of SLC38A5. Potential systemic influence of SLC38A5 knockout on circulating factors can be a compounding factor in interpretation of the results. Yet our data from ocular siRNA delivery strongly suggest that local inhibition of SLC38A5 did directly impair retinal angiogenesis, and systemic influence from other organs is likely minimal. It is also not clear whether other retinal cell types, including glia and neurons may be impacted by SLC38A5 modulation and thus affect vascular endothelium, although our cell culture data largely supports an EC-specific pro-angiogenic role of SLC38A5.

In summary, our data present direct evidence that SLC38A5 is a novel regulator of retinal angiogenesis. Expression of SLC38A5 is enriched in sprouting neovessels and driven directly by Wnt/β-catenin signaling pathway, as evident in Wnt-deficient retinas. Suppression of SLC38A5 may limit glutamine uptake by ECs, resulting in dampened VEGFR2 signaling and blunted retinal angiogenesis. Our findings of SLC38A5 as a modulator of pathologic retinal angiogenesis suggest the possibility of targeting SLC38A5 and its transported AAs as new therapeutic intervention for the treatment of vascular eye diseases and potentially other angiogenesis-related diseases.

## Acknowledgement

We thank Drs. Lois E. H. Smith, Ye Sun, and Bertan Cakir for very helpful discussion.

## Author contributions

Z.W. and J.C. conceived and designed the study; Z.W. and J.C. wrote the manuscript. Z.W., F.Y., S.H., C.-H.L., W.R.B., S. S.C., A.K.B., Y. T. and Z. F. performed experiments and collected and analyzed the data; Z. F., J.-X. M, and W.-H. L. shared reagents and resources and provided expert advice; all authors edited and approved the manuscript.

## Data and materials availability

This paper and the Supplementary Materials contain all data needed to evaluate the conclusions.

---

**Supplemental Figure S1.**
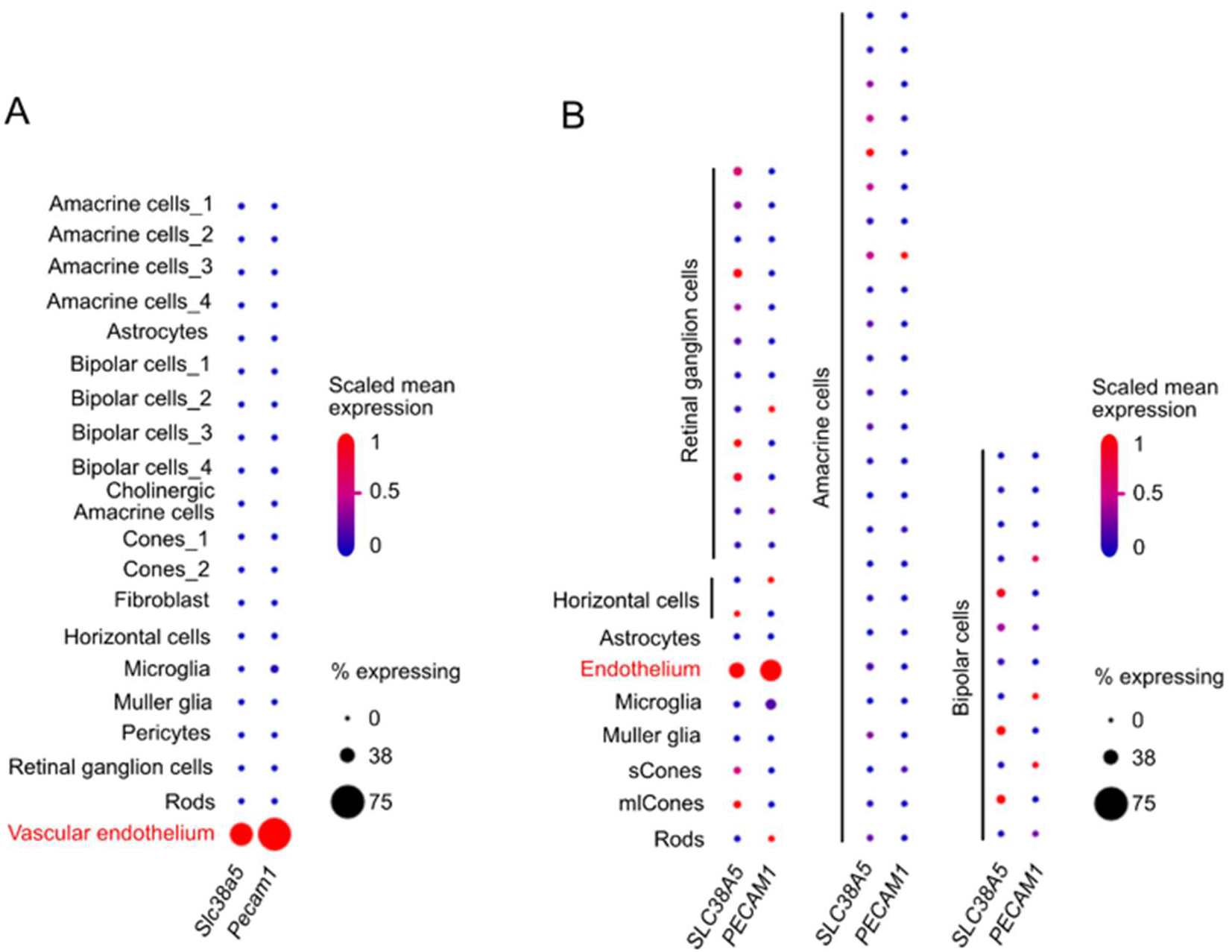
Distinct expression of *Slc38a5* in vascular endothelium in mouse and human retina with single-cell transcriptomics. **A**. Dot plot of *Slc38a5* and endothelial cell marker *Pecam1* gene expression (scaled) for different retinal cell types in P14 C57BL/6J mouse retinas. *Slc38a5* was distinctly expressed in vascular endothelium cluster. Data source: Study - P14 C57BL/6J mouse retinas (https://singlecell.broadinstitute.org/single_cell/study/SCP301) (56). **B**. Dot plot of *SLC38A5* and endothelial cell marker *PECAM1* gene expression (scaled) for different retinal cell types at human fovea and peripheral retina. *Slc38a5* was highly expressed in endothelium cluster. Data source: Study - Cell atlas of the human fovea and peripheral retina (https://singlecell.broadinstitute.org/single_cell/study/SCP839) (57). Scaling is relative to each gene’s expression across all cells of a cluster. Gene expression is scaled from zero to one (0.5 = the mean across all cells in the cluster file referenced for dot plotting). % expressing is the percent of cells that have one or more transcripts for the gene of interest.

**Supplemental Figure S2.**
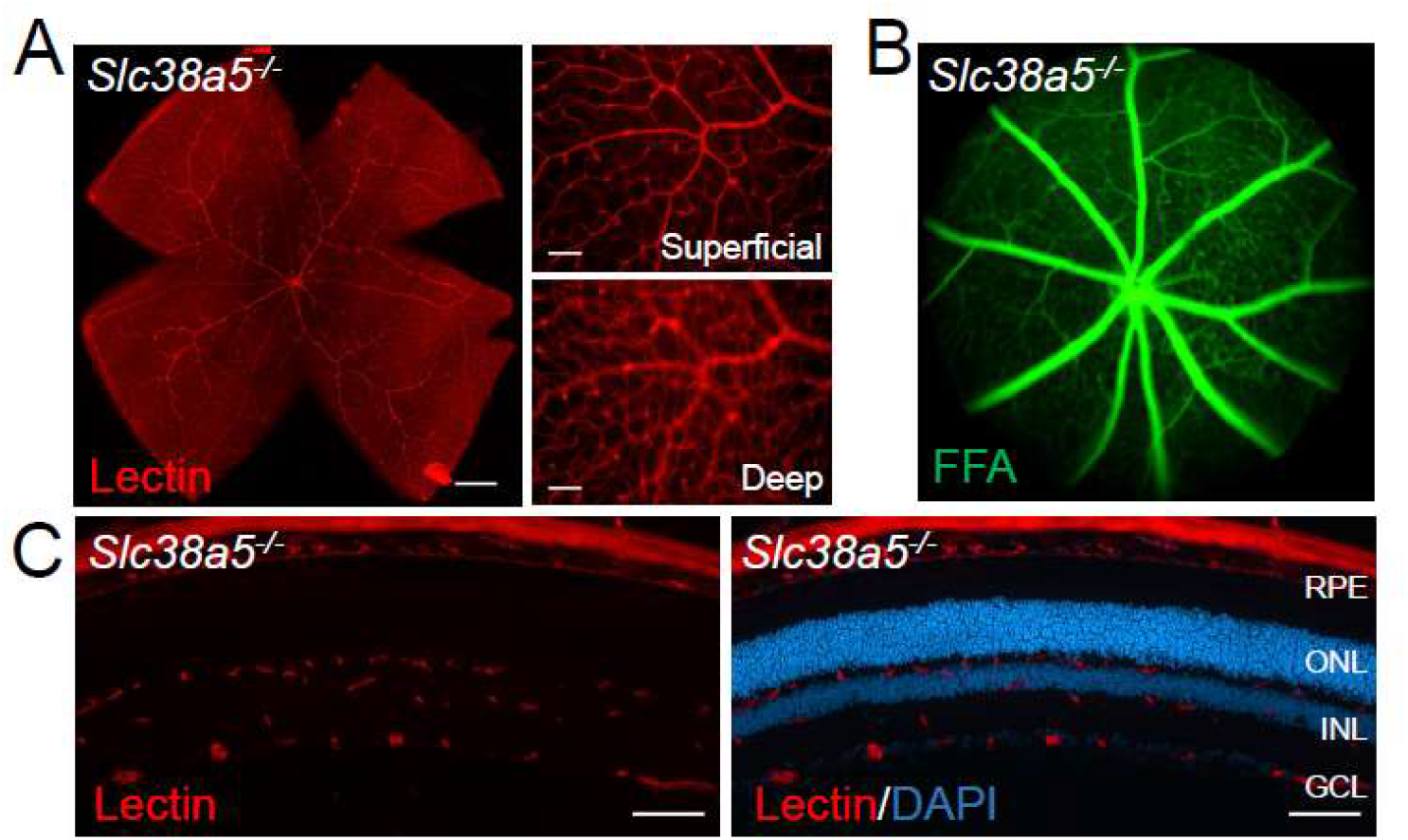
Adult *Slc38a5^-/-^* retinas have normal retinal blood vessels and vascular barrier. **A**: Flat mounts of adult WT and *Slc38a5^-/-^* retinas show normal branching and structure of retinal vessels stained with isolectin B4 (red). **B:** Fundus fluorescein angiography (FFA) of adult WT and *Slc38a5^-/-^* mice shows no sign of vascular leakage of fluorescein (green), indicating intact retinal vessels barrier in *Slc38a5^-/-^* eyes. **C:** Cross sections of *Slc38a5^-/-^* eyes show three normal layers of retinal vessels stained with isolectin B4 (red) and DAPI (blue). RPE: retinal pigment epithelium, ONL: outer nuclear layer, INL: inner nuclear layer, GCL: ganglion cell layer. Scale bars: A, left: 500 μm, right: 100 μm; C, 100 μm.

